# Using energy to go downhill – a genoprotective role for ATPase activity in DNA topoisomerase II

**DOI:** 10.1101/2023.06.27.546777

**Authors:** Afif F. Bandak, Tim R. Blower, Karin C. Nitiss, Viraj Shah, John L. Nitiss, James M. Berger

## Abstract

Type II topoisomerases effect topological changes in DNA by cutting a single duplex, passing a second duplex through the break, and resealing the broken strand in an ATP-coupled reaction. Curiously, most type II topoisomerases (topos II, IV, and VI) catalyze DNA transformations that are energetically favorable, such as the removal of superhelical strain; why ATP is required for such reactions is unknown. Here, using human topoisomerase II β (hTOP2β) as a model, we show that the ATPase domains of the enzyme are not required for DNA strand passage, but that their loss leads to increased DNA nicking and double strand break formation by the enzyme. The unstructured C-terminal domains (CTDs) of hTOP2β strongly potentiate strand passage activity in the absence of the ATPase regions, as do cleavage-prone mutations that confer hypersensitivity to the chemotherapeutic agent etoposide. The presence of either the CTD or the mutations lead ATPase-less enzymes to promote even greater levels of DNA cleava*in*ge*vitro*, as well as *in vivo*. By contrast, the aberrant cleavage phenotypes of these topo II variants is significantly repressed when the ATPase domains are restored. Our findings are consistent with the proposal that type II topoisomerases acquired an ATPase function to maintain high levels of catalytic activity while minimizing inappropriate DNA damage.

## Introduction

The action of DNA replication and transcription machineries promotes the supercoiling and entanglement of chromosomal DNA (Sundin and Varshavsky, 1980; Liu and Wang, 1987; Wang, 1994; Baranello *et al*., 2012). Cells resolve such topological challenges using a class of enzymes known as DNA topoisomerases (reviewed in (Chen, Chan and Hsieh, 2013)). Of all topoisomerases, the type II enzymes are distinguished by an ability to physically pass one DNA duplex through a transient, enzyme-mediated break in a second double-stranded DNA segment (Brown and Cozzarelli, 1981). In the type IIA topoisomerase subgroup, the class found most broadly across cellular organisms, strand passage is facilitated by the sequential opening and closing of three dissociable subunit-subunit interfaces – termed “gates” – in a manner controlled by ATP turnover (Wigley *et al*., 1991; J Roca and Wang, 1994; Berger *et al*., 1996; Roca *et al*., 1996a; Morais Cabral *et al*., 1997). DNA gyrase, an archetypal prokaryotic type IIA topoisomerase, consumes ATP to power the introduction of negative supercoils into DNA, an energy-requiring reaction (Gellert *et al*., 1976). Interestingly, gyrase also has been shown to be capable of catalyzing ATP-in dependent supercoil relaxation (Gellert *et al*., 1977; Brown, Peebles and Cozzarelli, 1979); however, while this activity does not require the presence of its ATPase elements, it does depend on its specialized C-terminal DNA wrapping domains (Reece and Maxwell, 1991). By comparison, other type IIA topoisomerases such as eukaryotic topo II and prokaryotic topo IV have been thought to strictly require ATP to perform supercoil relaxation (Goto and Wang, 1982; Peng and Marians, 1993), even though this reaction is energetically favorable.

Why non-supercoiling type IIA topoisomerases consume ATP when product formation holds no energetic cost has been a long-standing question (Bates and Maxwell, 2010). Evolutionarily, the DNA binding and cleavage element of type IIA family members is thought to have been augmented with a GHKL-family ATPase domain to give rise to modern-day type IIA topoisomerases (Forterre *et al*., 2007). It has been suggested that the widespread success of these ATP-dependent type II topoisomerases emerged from an evolutionary pressure to use nucleotide turnover as a mechanism to control conformational changes that regulate DNA cleavage and guard against the accidental formation of DNA breaks (Bates, Berger and Maxwell, 2011). Although structural and biochemical studies have provided evidence that the ATPase cycle is indeed coupled to large-scale physical rearrangements in type II topoisomerases (Osheroff, Zechiedrich and Gale, 1991), the idea that it can also regulate DNA break formation to promote DNA integrity has lacked experimental support.

To better understand the role of ATP in controlling type IIA topoisomerase activity, we constructed and analyzed the biochemical and cellular activities of several truncations and hyper-cleavage mutants of human topoisomerase IIβ (hTOP2β), one of two type IIA topoisomerase isoforms found in human cells (Austin and Marsh, 1998). We show that the hTOP2β nucleolytic core (which lacks both the N-terminal ATPase domain and a poorly conserved, intrinsically disordered C-terminal region) is capable of supercoil relaxation in the absence of ATP, albeit at relatively low levels compared to the ATP-stimulated activity of the full-length enzyme. In the presence of the C-terminal domain (CTD), the ATP-independent topoisomerase activity of the core is boosted substantially, to levels approaching that of full-length hTOP2β with ATP, but at a cost of elevated DNA damage propensity. Interestingly, we recently showed that certain point mutations in full-length hTOP2β, such as hTOP2β^R757W^(Bandak *et al*., 2023), would modestly increase persistent DNA cleavage events *in vitro* and elicit moderate hyper-sensitivity to the topo II poison etoposide *in vivo*. These mutations also boosted the ATP-independent activity of the nucleolytic core to levels comparable to that o f full-length h TOP2β but w ith a m arked increase i n aberrant DNA s trand breakage ((Bandak *et al*., 2023)). By contrast, restoration of the ATPase domain to either the core with the CTD or the R757W mutant substantially mitigates the hyper-cleavage phenotypes of these enzyme variants. Together, our findings establish that an isolated type IIA topoisomerase DNA binding and cleavage core can indeed function as an ATP-independent strand passage enzyme and that this family of enzymes likely acquired its ATPase domains to minimize inappropriate DNA scission.

## Results

### The nucleolytic core of human hTOP2β possesses ATP-<u>in</u>dependent strand passage activity

To begin to define the relative contributions of individual type IIA topoisomerase domains to overall enzyme function, we first characterized the activities of various truncations of hTOP2β, starting with the DNA binding and cleavage domains of the enzyme (the nucleolytic core, residues 447-1206). Increasing amounts of the hTOP2β core were incubated with negatively supercoiled plasmid DNA in the absence of ATP for 30 min; reactions containing full-length hTOP2β were run in parallel both with and without ATP (**Fig. 1A**). Reactions were quenched with SDS and proteinase K and analyzed using native agarose gel electrophoresis. As expected, full-length hTOP2β showed robust supercoil relaxation activity in the presence of ATP (fully relaxing 250 ng of plasmid in 30 min at an enzyme dimer:DNA ratio of ∼2.5:1) and no relaxation was observed with oFuigt .A1TAP).(By contrast, a low level of relaxation of the supercoiled DNA substrate was evident as the concentration of the hTOP2β core was increased to 10:1 or 20:1 protein dimer:plasmid (**Fig. 1A**). Because the core lacks the ATPase domains and no nucleotide was present in the reaction buffer, the observed activity is clearly ATP-<u>in</u>dependent.

**Fig. 1.**
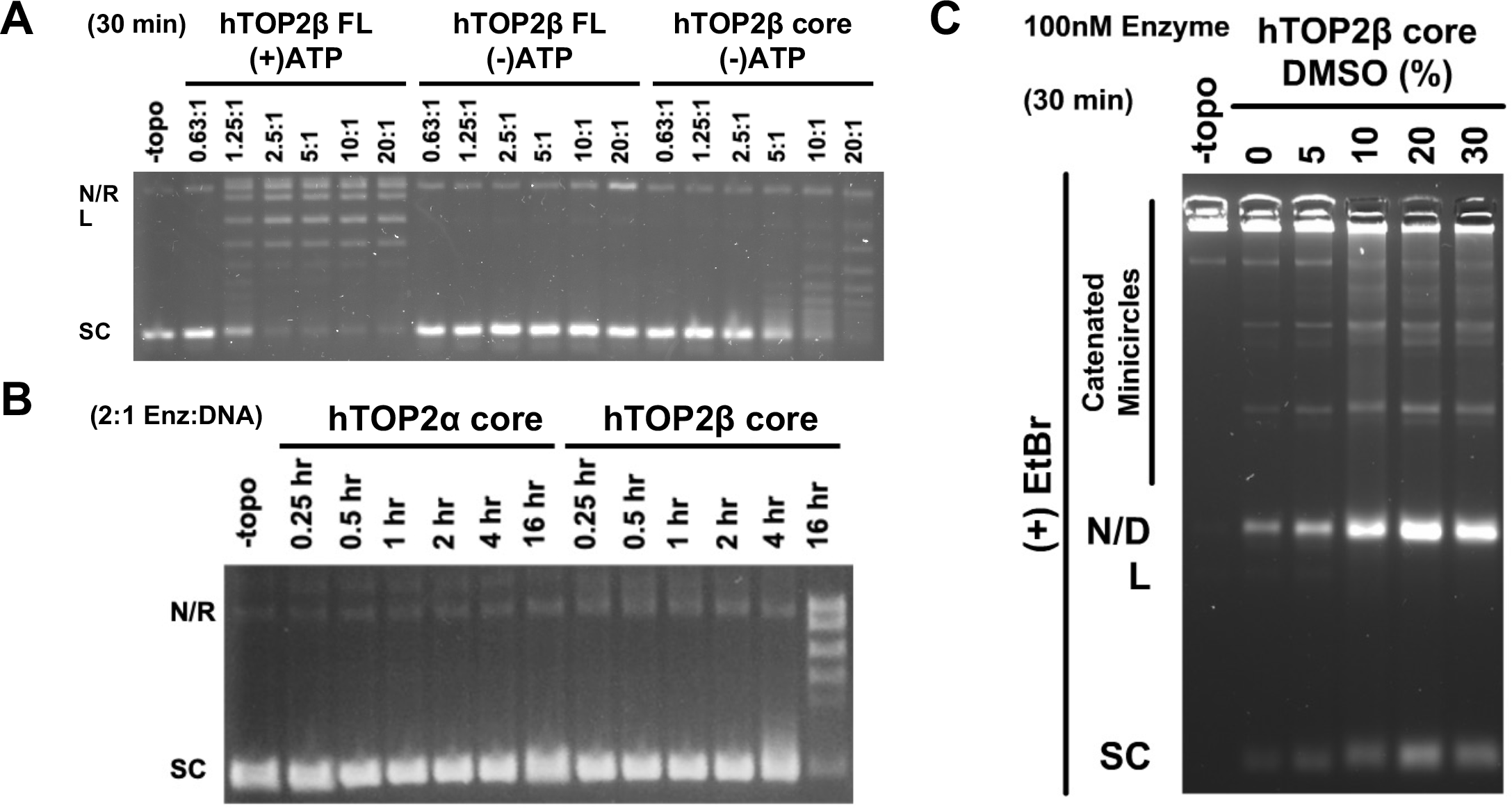
The hTOP2β core can perform supercoil relaxation and decatenation. (A) Activity of core and full-length hTOP2β enzymes on negatively supercoiled plasmid DNA in the presence (+) or absence () of ATP. Ratios refer to enzyme dimers:DNA. (B) Time course of hTOP2α and hTOP2β core enzymes acting on negatively supercoiled plasmid DNA. (C) Decatenation of kDNA by the hTOP2β core as titrated against different concentrations of DMSO. Enzyme (100 nM) was incubated with 250 ng of kDNA for 30 min at 37 °C.

To further investigate the ATP-independent supercoil relaxation activity of the hTOP2β core, we compared its functionality with the nucleolytic core of another topo II enzyme, human topoisomerase IIα (hTOP2α, residues 431-1193). Native agarose gel electrophoresis of the reactions showed that at a near stoichiometric enzyme-to-DNA ratio (2:1), the hTβOPco2re could relax nearly all the supercoiled plasmid substrate, but slowly, over the course of ∼16 h (**Fig. 1B**). By comparison, hTOP2α failed to generate detectable relaxed products over the same time course (**Fig. 1B**). We next examined reactions over a shorter time-frame (30 min) but using a higher protein concentration (20:1 dimer:plasmid) (Fig. S1A). Different concentrations of DMSO (a mild protein destabilizing and drug solubilizing agent (Tjernberg, Markova, Griffiths and HalléN, 2006)) were also included in this assay to determine whether the reagent might perturb the interactions between subunit interface and thereby enhance relaxation activity. Under these conditions, the hTOP2β core could be seen to relax supercoiled DNA even in the absence of DMSO, and its activity improved substantially as DMSO concentrations were increased (Fig. S1A). Interestingly, the hTOP2α core was also able to carry out a limited degree ATP-independent supercoil relaxation activity when DMSO concentrations were increased, although to a lesser extent than the hTOP2β nucleolytic core (Fig. S1A). To confirm that the supercoil-relaxation activity observed for the hTOP2β (and hTOP2α) core was not due to a contaminating topoisomerase in our protein preparations, we purified mutant forms of the proteins lacking the catalytic tyrosine required for DNA cleavage (hTOP2α^Y805F^ and hTOP2β^Y821F^) and assessed their activity on negatively supercoiled DNA substrates (Fig. S1B). Neither mutant protein showed any ability to relax, cleave, or nick the substrate DNA, regardless of DMSO concentration, demonstrating that the supercoil-relaxation activity seen with preparations of the wildtype cores derives from the purified human TOP2s (Fig. S1B). Collectively, this analysis shows that the cores of different type IIA topoisomerases can act in an ATP-independent manner, but with distinct innate efficiencies.

During our studies, it occurred to us that the supercoil relaxation activity we observed might be due to some type of non-canonical nicking and religation reaction, rather than due to strand passage. To distinguish between these possibilities, we assessed whether the hTOP2β core could decatenate a kDNA substrate. kDNA consists of a network of catenated DNA circles that cannot enter the wells of an agarose gel unless liberated by type II topoisomerases (Marini, Miller and Englund, 1980). As with the supercoil relaxation assay, the hTOP2β core displayed a weak ability to produce decatenated circles; however, the efficiency of this reaction increased markedly with increasing DMSO concentration (**Fig. 1C**). These data demonstrate that the central DNA binding and cleavage region of hTOP2β is indeed sufficient to carry out a *bone fide* strand passage event.

### Strand passage by the hTOP2β core can be markedly increased by the C-terminal domain or a cleavage-prone point mutation

After discovering that the hTOP2β nucleolytic core is capable of ATP-independent strand passage, we next sought to determine whether the C-terminal domain of the enzyme has any effect on this activity. The C-terminal regions of eukaryotic type IIA topoisomerases generally consist of long, unstructured elements (>200 amino acids) that are poorly conserved between orthologs (Linka *et al*., 2007). We generated a ‘headless’ hTOP2β (hTO Pβ2-HL) construct that includes all but the ATPase domain of the enzyme and assessed its ability to relax negatively supercoiled DNA compared to both full-length hTOP2β and the hTOP2β catalytic core on native agarose gels. In parallel, reaction products were also examined on agarose gels in the presence of ethidium bromide to assess whether the ATPase-less proteins exhibited any elevated propensity to nick or cleave DNA.

Surprisingly, the hTOP2β-HL construct proved capable of relaxing supercoiled DNA in the absence of ATP more efficiently than the hTOP2β core (∼4-fold, as evidenced by the amount of enzyme required to deplete a majority of the starting supercoiled substate over a 30 min reaction period) (**Fig. 2A**). Indeed, the ATP-independent supercoil relaxation activity of the hTOP2β-HL construct was now decreased by only ∼4-fold compared to the ATP-dependent relaxation activity of full-length hTOP2β, although it did appear substantially less processive (as evidenced by the continuous ladder of partially relaxed plasmid topoisomers that were produced) (**Fig. 2A**). An analysis of gels run with ethidium bromide showed that hTOP2β-HL also produced higher levels of nicked and linearized plasmid species compared to wildtype hTOP2β under conditions where total cleavage activity (both reversible and irreversible, observed by using an SDS-only quench) was assessed (**Fig. 2B**). By comparison, when only irreversible cleavage was monitored (by adding EDTA to the reactions prior to the addition of SDS), a more modest degree of elevated nicking was seen for the core, accompanied by a still higher level of both nicking and linearization for the headless construct (**Fig. 2C**). Overall, the removal of the type IIA topoisomerase ATPase domains correlates with an increased propensity for the enzyme to generate DNA cleavage products, with constructs exhibiting higher strand passage activity also proving more likely to generate dsDNA breaks.

**Fig. 2.**
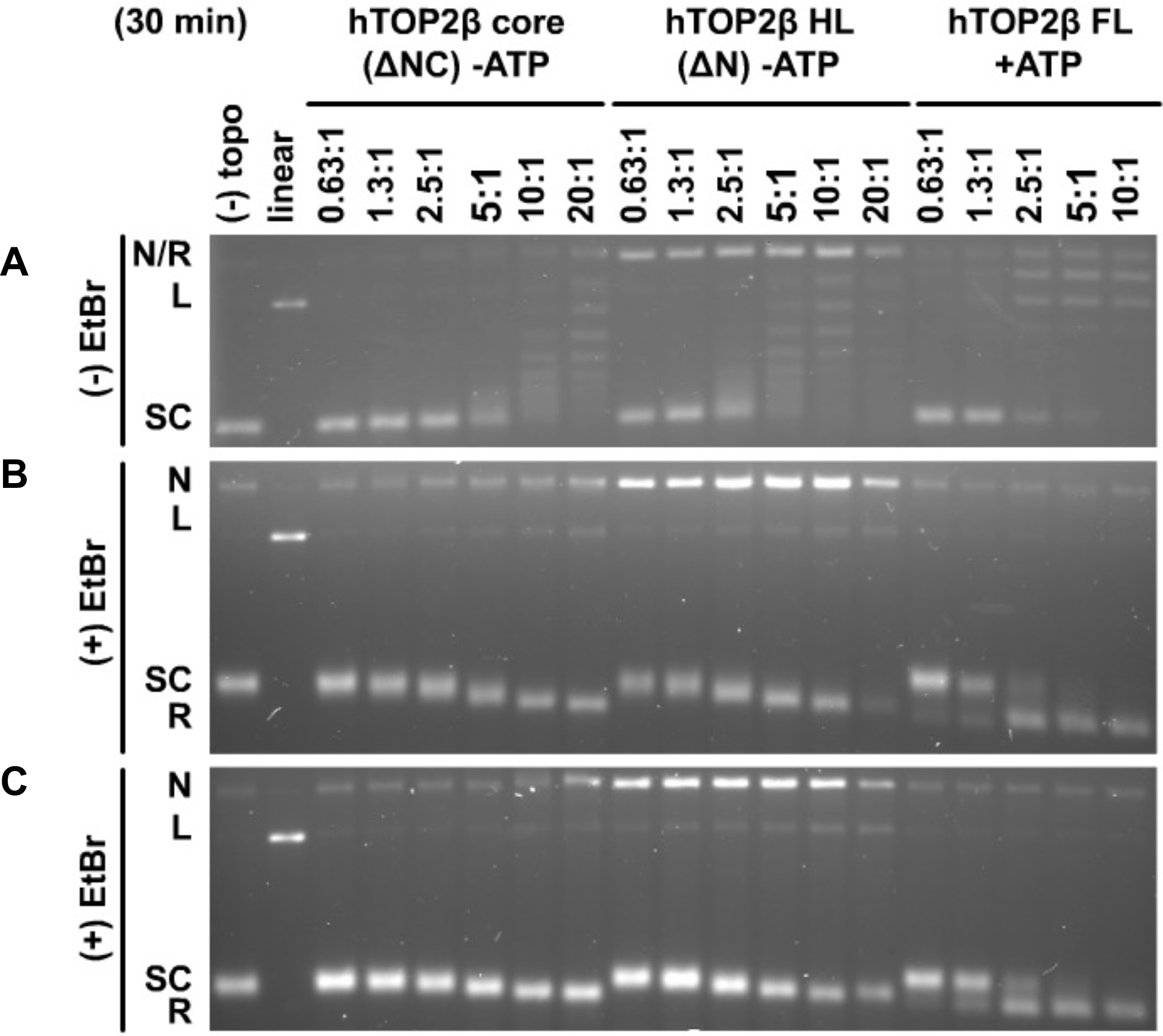
Headless hTOP2β has enhanced activity over the hTOP2β core but is more cleavage prone than full-length hTOP2β. (A) Relaxation activity of the hTOP2β core, headless hTOP2β (HL) and full-length (FL) hTOP2β on negatively supercoiled plasmid DNA in the presence (+) or absence (-) of ATP. Ratios refer to enzyme dimers:DNA. (B and C) Assays as per (A) but with products separated on agarose gels containing ethidium bromide (EtBr), having either been quenched only with SDS (B, irreversible and reversible products) or with EDTA followed by SDS (C, irreversible products only).

While conducting these studies, a parallel effort in our groups identified a single amino acid substitution in hTOP2β, R757W, which could strongly potentiate the sensitivity of the enzyme to the anti-cancer agent etoposide, but at a cost of elevating its innate DNA cleavage activity(Bandak *et al*., 2023). Interestingly, Arg757 forms part of the dimer interface that comprises the DNA-binding and cleavage site of hTOP2β (the DNA-gate) (Fig. S2A). This juxtaposition, coupled with the mutant’s hyper-cleavage phenotype, led us to hypothesize that replacing Arg757 with Trp might destabilize the interface, making it more prone to spontaneous separation. To test this hypothesis, we examined whether the R757W substitution could stimulate the ATP-independent supercoil relaxation activity of the hTOP2β core. Enzyme titrations showed that the R757W core mutant was highly active relative to the wildtype hTOP2β core (∼20-fold greater,**Fig. 3A**), producing fully relaxed product over 30 min at a roughly 1:1 protein dimer:DNA ratio (a level of activity comparable to that of full-length hTOP2β with ATP, see **Fig. 1A**). The R757W mutant was similarly more active in kDNA decatenation than the wildtype hTOP2β core (down only 8-fold compared to full-length enzyme), even in the absence of DMSO (Fig. S2B). To determine if this phenomenon is limited to the R757W variant, we tested whether another known hyper-cleavage mutation (K600T,(Bandak *et al*., 2023)) could stimulate ATP-independent supercoil relaxation activity. Enzyme titrations similarly showed that the K600T core mutant was highly active relative to the wildtype hTOP2β core (Fig. S3).

**Fig. 3.**
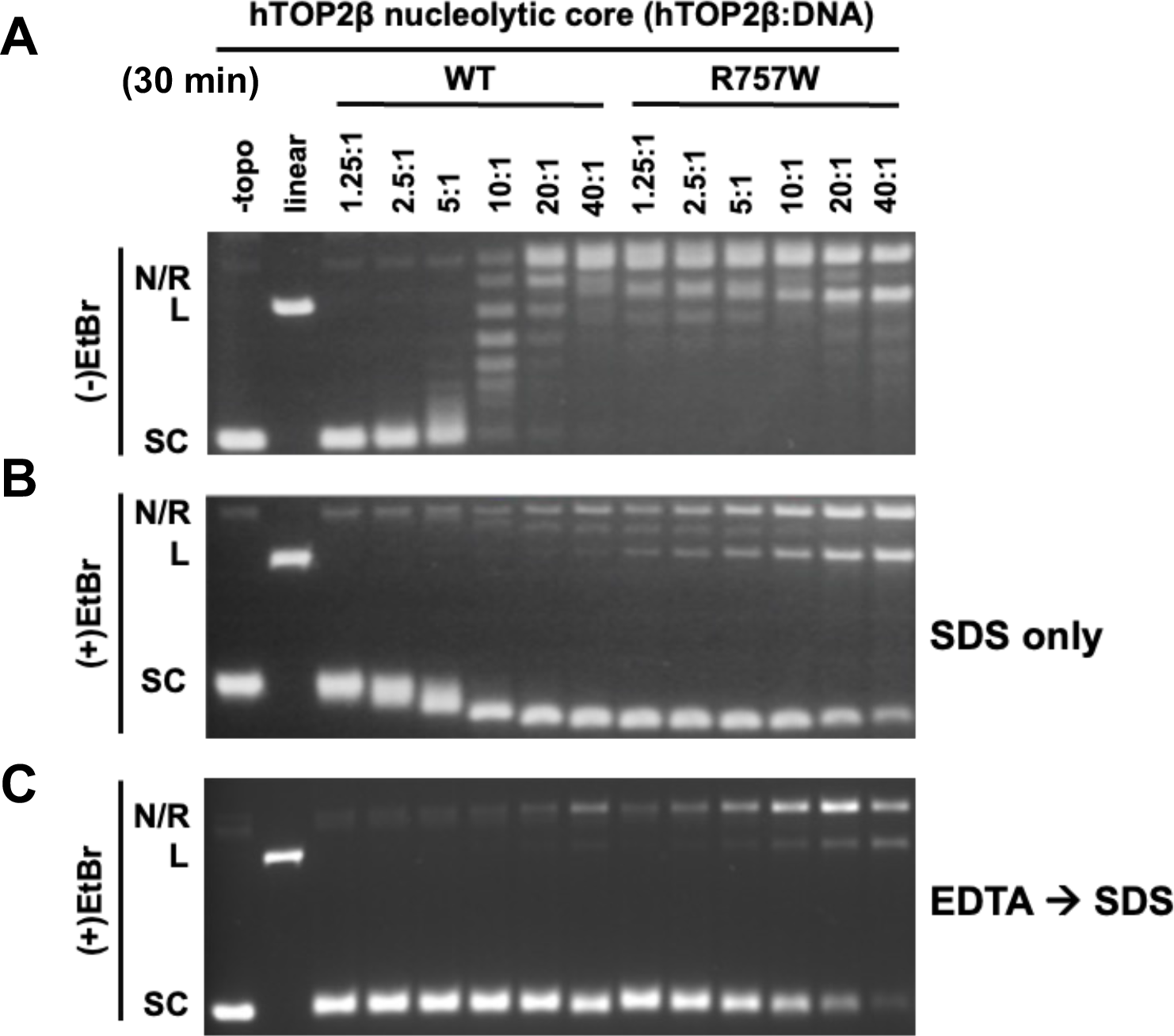
The hTOP2β^R757W^ core has enhanced DNA supercoil relaxation activity, as well as enhanced reversible and irreversible DNA nicking and cleavage activity. (A) Relaxation activity of the wildtype hTOP2β and hTOP2β^R757W^ cores on negatively supercoiled plasmid DNA. Ratios refer to enzyme dimers:DNA. (B and C) Assays as per (A) but with products separated on agarose gels containing ethidium bromide (EtBr), having either been quenched only with SDS (B, irreversible and reversible products) or with EDTA followed by SDS (C, irreversible products only).

Reactions were also analyzed by agarose gels run in the presence of ethidium bromide in order to assess the impact of the Arg→Trp mutation on DNA nicking and linearization, (**Fig. 3B-C**). Here, the hTOP2β^R757W^ core construct displayed significantly elevated levels of both reversible and irreversible DNA cleavage products (**Fig. 3B-C**), even higher than the hTOP2β headless construct (**Fig. 2B-C**). However, when the R757W mutation was reintroduced back into full-length hTOP2β, it showed considerably lower levels of nicked and linear product formation (Fig. S4). Collectively, these findings establish that the type IIA topoisomerase core can be converted into a robust, ATP-independent strand passage enzyme by the addition of the CTDs or even a single amino acid substitution, but that this elevated activity is typically accompanied by increased DNA cleavage activity. These observations also show that DNA damage-promoting alterations in hTOP2β can in turn be markedly attenuated by the presence of the ATPase domains.

### Full-length hTOP2β can also perform ATP-independent relaxation

Given the activities of the core and headless hTOP2β constructs, we next sought to examine whether full-length hTOP2β might have some capacity to relax DNA in an A TP-independent manner Reasoning that such an activity might be slow and/or inefficient, we initially tested a range of reaction incubation times, from 30 minutes to 8 hours. As compared to studies of the topo II cores that were conducted at a high protein dimer:DNA ratio (20:1) – which allowed activity to be seen on a short (30 min) time scale – here, we used a lower protein concentration (4:1 dimer:DNA). When tested from 30 min to 8 hours at this enzyme concentration, full-length hTOP2β displayed a barely detectable capacity to relax supercoiled DNA in the absence of ATP (**Fig. 4A**). As a comparison, full-length hTOP2α did not exhibit any relaxation activity, but full activity for both hTOP2α and hTOP2β was seen if ATP was added at the 8h mark, indicating the proteins were still active (**Fig. 4A**). Interestingly, when assayed in the context of the R757W mutation, the full-length hTOP2β mutant proved relatively robust at removing negative supercoils in an ATP-independent manner (**Fig. 4A**). At longer reaction times (16 h) and even lower protein dimer:DNA ratios (2.5:1), ATP-independent supercoil relaxation by full-length, wildtype hTOP2β was also clearly evident and, as seen for the hTOP2β core, was further stimulated by DMSO (supercoil relaxation activity was again not observed for full-length hTOP2α, even in the presence of increasing DMSO concentrations) (**Fig. 4B**). In all cases examined, ATP-independent supercoil relaxation was abolished by EDTA, indicating that the activity we observed is attributable to type IIA topoisomerase activity and not to a contaminating type IB topoisomerase, which carries out relaxation in the absence of a divalent cation (**Fig. 4A, hTOP2β^R757W^ final lane and Fig. 4B**).

**Fig. 4.**
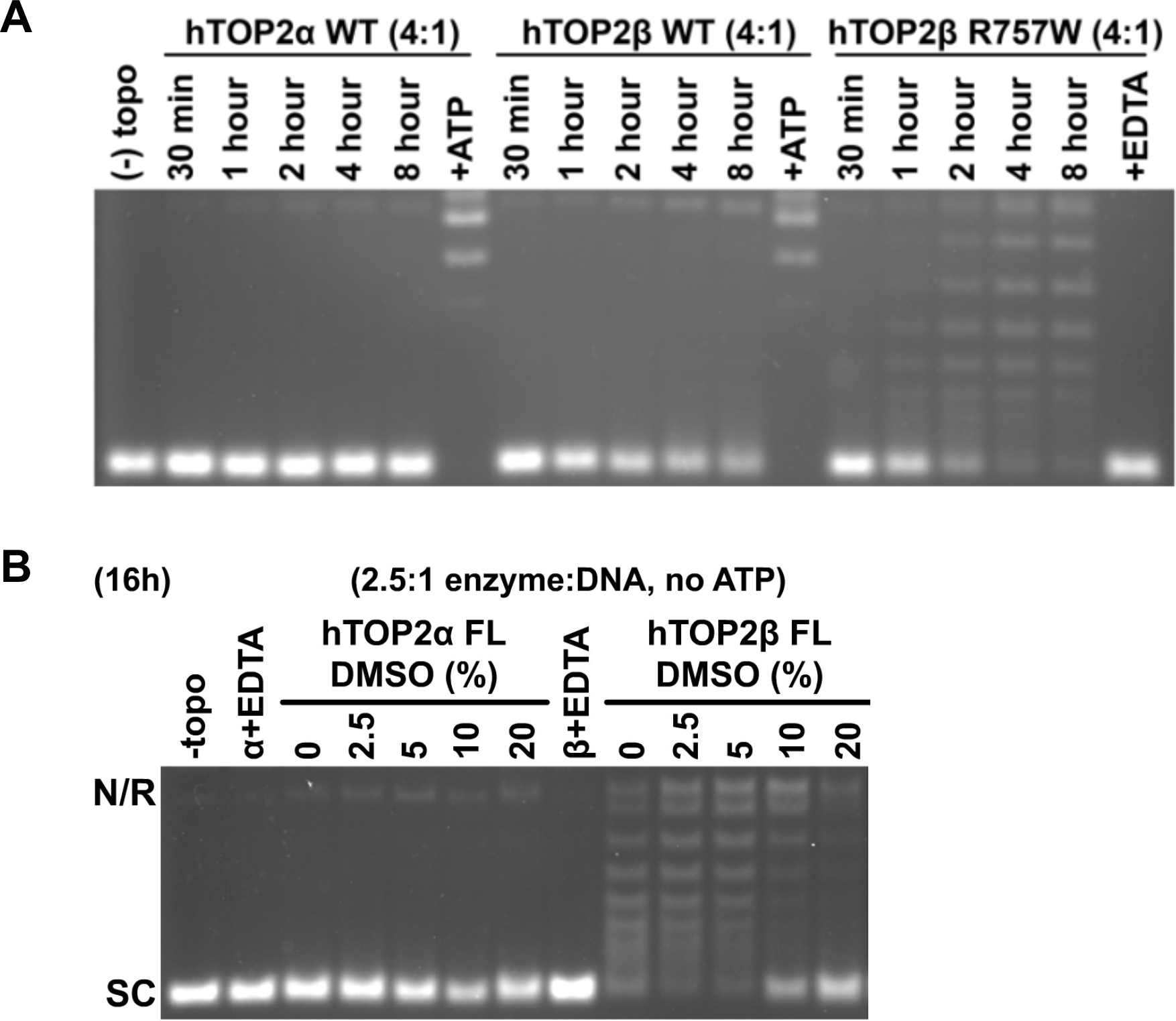
Full-length hTOP2β relaxation activity in the absence of ATP. (A) Time course of relaxation activity for wildtype, full-length hTOP2α and hTOP2β and full-length hTOP2β^R757W^ in the absence of ATP. Ratios refer to enzyme dimers:DNA. Single-lane controls were performed either in the presence of ATP (hTOP2α and hTOP2β) or EDTA (hTOP2β^R757W^). (B) Relaxation activity for wildtype, full-length hTOP2α and hTOP2β titrated against DMSO in the absence of ATP. Single-lane controls were performed for both enzymes in the presence of EDTA.

### ATPase-less hTOP2β does not complement a top2-4 temperature sensitive yeast mutation but generates DNA damage in yeast cells

Because headless hTOP2β exhibited relatively robust strand-passage activity compared to the full-length enzyme (**Fig. 2A**), we next asked whether this construct could support any of the essential functions of topo II in budding yeast. The nuclear localization signal of full-length hTOP2β resides in its C-terminal domain (Mirski, Gerlach and Cole, 1999), which allowed us to assess the cellular potential of hTOP2β-HL but not the hTOP2β core, which lacks this element. A plasmid expressing the hTOP2β headless construct under the control of the yeast *TPI1* promoter was introduced into *top2-4* temperature sensitive cells that also contain a wildtype allele of the *RAD52* gene, which encodes a protein important for the repair of DNA double-strand breaks. The expression of hTOP2β-HL did not affect cell survival at 25 °C but failed to complement growth at the non-permissive temperature of 34 °C (**Fig. 5A**). Because the R757W mutant of the hTOP2β-core exhibited even greater strand-passage activity than the wildtype core (**Fig. 3**), we also constructed a headless version of this mutant (hTOP2β-HL ^R757W^). Biochemical studies confirmed that, as with the core alone, the R757W mutation also substantially simulated the DNA supercoil relaxation and DNA decatenation activities of the headless construct compared to wildtype hTOP2β-HL, reaching levels matching that of purified, full-length hTOP2β (Fig. S5). However, when transformed into *RAD52+ top2-4* cells, hTOP2β-HL ^R757W^ also failed to complement the growth at the non-permissive temperature (**Fig. 5A**). Thus, despite the robust DNA strand passage activity displayed by both the headless and R757W mutant enzymes, hTOP2β still requires its ATPase domains to support its essential functions in yeast cells.

**Fig. 5.**
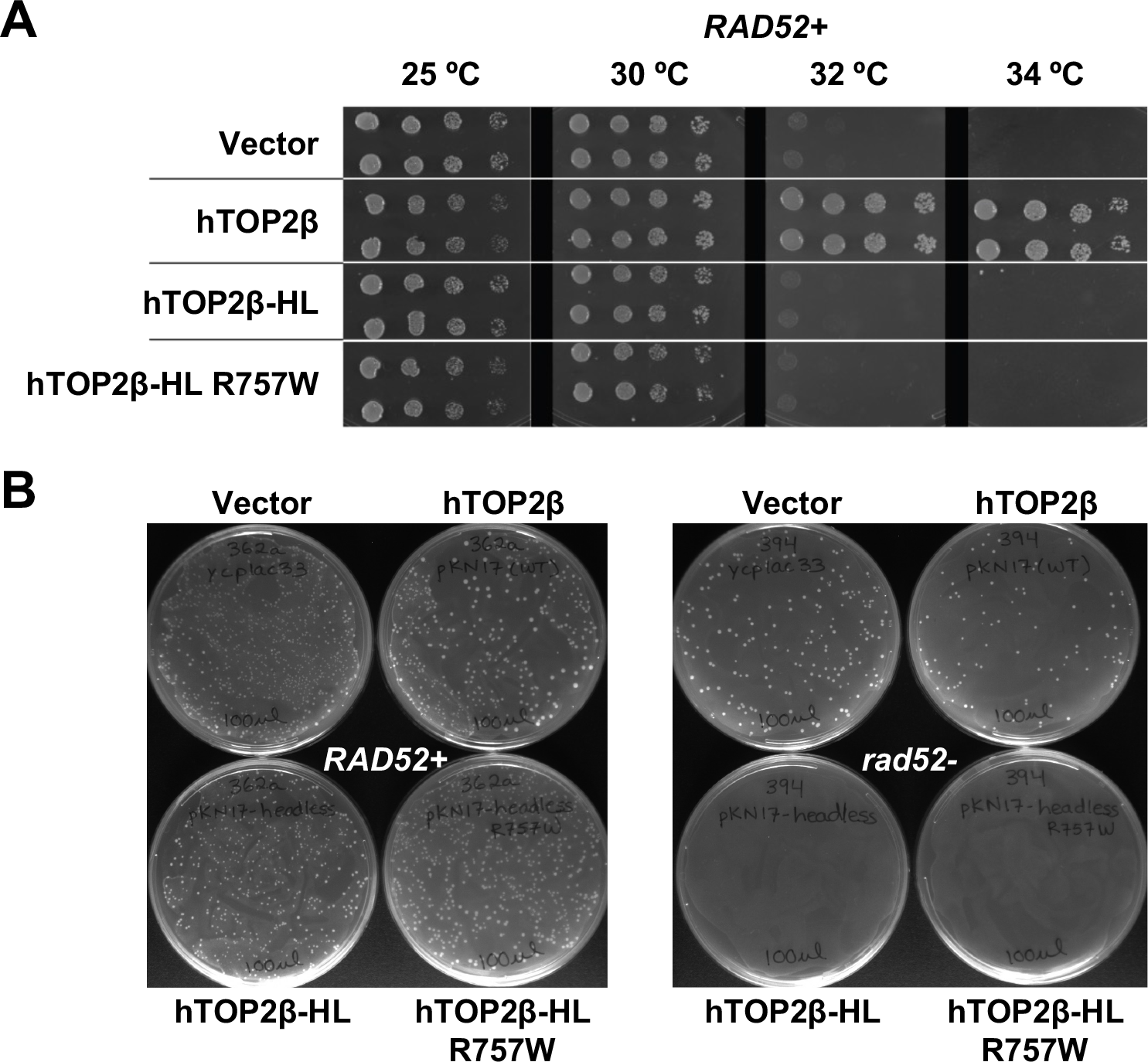
Headless hTOP2β fails to complement yeast growth and is toxic to repair-deficient cells. (A) Temperature-dependent complementation of *top2-4* yeast strain by hTOP2β wildtype, full-length (FL) and headless (HL) hTOP2β and headless hTOP2β^R757W^. (B) Transformants obtained upon complementation of *TOP2*+ yeast cells with hTOP2β constructs, in either *RAD52+* or *rad52-*backgrounds.

Because we found that ATPase-less topo II constructs innately generate more cleavage products than full-length hTOP2β *in vitro*, we next tested whether this damage propensity might also manifest *in vivo*. Upon introducing either hTOP2β-HL or hTOP2β-HL ^R757W^ into a yeast background proficient for native topo II activity but deficient in Rad52 function (*TOP2+ rad52-*), we found we could obtain no colonies (**Fig. 5B**). This result establishes that the ATPase-less hTOPβ2 constructs are not able to complement normal cellular growth but are also actively generating DNA damage. To further test this idea, hTOP2β-HL was introduced into a diploid yeast strain, CG2009, which carries several heteroallelic markers for assessing homologous recombination(Stantial *et al*., 2020). We then measured recombination frequencies in CG2009 cells carrying an empty vector, full-length hTOP2β, or hTOP2β-HL. Significantly, cells transformed with hTOP2β-HL displayed an approximately 50-fold elevation in recombination frequency compared to cells transformed with the parent plasmid expressing full-length hTOP2β (**Table S1**). Taken together, these results demonstrate that the DNA binding and cleavage core of hTOP2β is a potent DNA-damaging agent whose genome-destabilizing potential is masked by its ATPase elements.

## Discussion

Type II topoisomerases catalyze DNA strand passage events by generating transient double-strand breaks in chromosomal segments. This activity is both indispensable for and potentially detrimental to genetic integrity in cells (Wang, Caron and Kim, 1990); indeed, certain classes of clinically used drugs known as topoisomerase poisons can disrupt the enzyme’s DNA breakage-rejoining cycle to induce DNA damage and kill cells (Pommier et al., 2016). Here, we probed how specific domains of hTOP2β modulate the potential to form cleaved and uncleaved DNA states of the enzyme.

One of the more significant and unexpected observations for hTOP2β was the ability of its DNA binding and cleavage core – and even the full-length enzyme – to catalyze supercoil relaxation in the absence of AT PFi(gs. 1, 4), a nucleotide cofactor long thought to be essential for eukaryotic type II topoisomerase activity. Supercoil relaxation was not observed for the hTOP2α core F(ig. 1), although strand passage by the hTOP2α core (and by the equivalent region of hTOP2β) was stimulated by DMSO (Fig. S1), an agent commonly used to solubilize small-molecule drugs that is known to also have a mild destabilizing effect on proteins (Tjernberg, Markova, Griffiths and Hallén, 2006; Arakawa, Kita and Timasheff, 2007; Chan *et al*., 2017). DMSO was not nearly as efficient at promoting DNA supercoil relaxation by the hTOP2α core as compared to hTOP2β. These findings indicate that the dimer interfaces of hTOP2β can transiently separate to allow DNA strand passage. They also demonstrate that the interfacial stability of hTOP2β innately differs from its human paralog, hTOP2α, allowing for more promiscuous (i.e., nucleotide-uncoupled) DNA cleavage, supercoil relaxation, and decatenation activities. Although supercoil relaxation by the hTOP2β core appears somewhat more efficient than decatenation, the observation that this region can separate closed circles from a kDNA network confirms that the protein is performing DNA strand passage, as opposed to using a nick-and-swivel mechanism analogous to that employed by type IB topoisomerases. We speculate that the torsional energy stored in supercoiled DNA may help facilitate core activity by more readily promoting crossover formation. Although it is possible that the DNA binding and cleavage core could work by a “one-gate” approach in which the C-terminal dimer interface of the enzyme remains closed during strand passage, the simplest interpretation of the data is that the enzyme operates by the “two-gate” mechanism utilized for the full-length enzyme, wherein a transported DNA moves through the sequential opening and closing of the cleavage DNA segment (at the DNA-gate) and the C-terminal dimer interface (the ‘C-gate’) (J Roca and Wang, 1994; Roca *et al*., 1996a) (**Fig. 6A**).

**Fig. 6.**
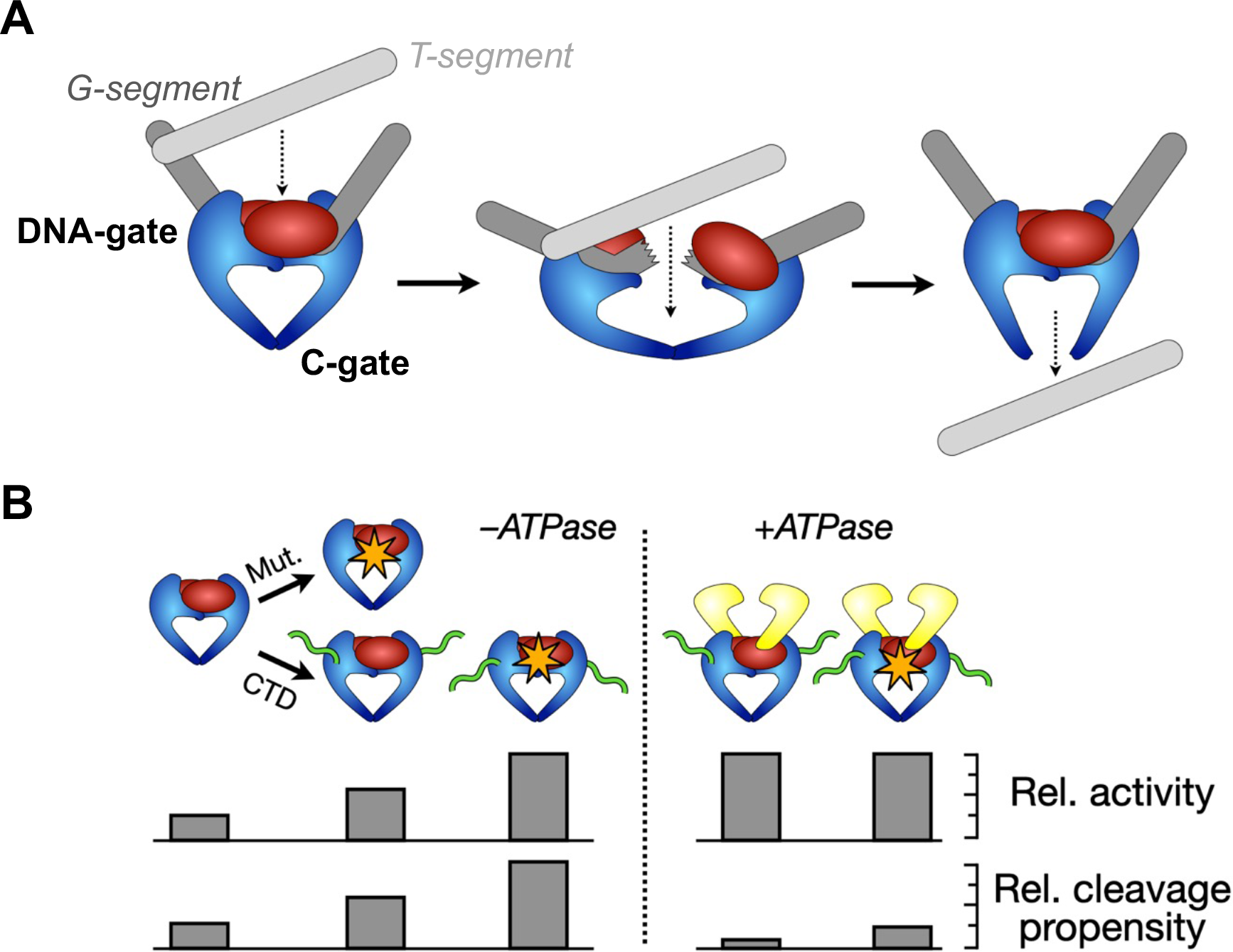
Schematic depicting the evolution of a hypothetical, damage-prone ancestral type II topoisomerase before the acquisition of a regulating ATPase element for mitigating unwarranted DNA breakage. (A) An ancestral, ATPase-less type II topoisomerase could perform strand passage by a “two-gate” mechanism that relies on just the DNA- and C-gates of the enzyme. Data reported here for wild-type and mutant (R757W) hTOP2β show that this minimal type II topoisomerase construct indeed possesses such an activity. (B) Acquisition of a C-terminal domain (CTD) and/or mutations that destabilize subunit interfaces can potentially lead to enhanced activity but also to an enhanced propensity for cleavage. The acquisition of the GHKL family ATPase domains substantially reduces deleterious DNA cleavage activity. Our findings suggest that the cleavage-prone behavior observed for hTOP2β in certain cellular contexts (such as transcriptional bursting (Ju *et al*., 2006; Haffner *et al*., 2010; Trotter, King and Archer, 2015)) may be the consequence of an inherent subunit interface lability retained from an ancestral enzyme, rather than from a cellular desire to generate DNA breaks.

The strand passage activity of the hTOP2β core is notable, as the only other type IIA topoisomerases currently known to be capable of acting in an ATP-independent manner are *E. coli* DNA gyrase and DNA topoisomerase II from bacteriophage T4 (Sugino *et al*., 1977; Liu, Liu and Alberts, 1980; Papillon *et al*., 2013). In gyrase, this function depends on a specialized DNA-wrapping domain that is not present in eukaryotic type IIA topoisomerases (Kampranis and Maxwell, 1996; Corbett, Shultzaberger and Berger, 2004). The DNA-binding and cleavage core of type IIA topoisomerases in general has been suggested to arise from diverse (likely viral) origins (Forterre and Gadelle, 2009), and the cores are thought to have acquired ATPase domains to regulate their activity. The hTOP2β core exemplifies the activity predicted for an ancient type IIA topoisomerase (Bates, Berger and Maxwell, 2011), one that arose from a modified type of nuclease, before the acquisition of a modulatory ATPase element (**Fig. 6A**). Interestingly, we found that the unstructured C-terminal region of hTOP2β stimulates the ATP-independent activity of core (**Fig. 2**), establishing that it plays a role in supporting DNA strand passage. The C-terminal tail of eukaryotic topo II has been recently shown to promote protein-DNA interactions that can modulate the catalytic output of the enzyme (Jeong *et al*., 2022). It seems likely that this element boosts the activity of the hTOP2β core through these interactions, although how this potentiation occurs at the molecular level has yet to be established.

It has been suggested that the pressure to reduce DNA damage arising from unregulated DNA cleavage events could have been an evolutionary driving force towards the coupling of strand passage to ATP turnover in type II topoisomerases (Bates, Berger and Maxwell, 2011). In this view, ATPase activity would serve both to promote the sequential separation of subunit interfaces in the core (which could now be strengthened by natural selection to avoid spontaneous opening) and as a switching mechanism to ensure that cleaved DNA is resealed before the ATPase domain dimer interface (termed the ‘N-gate’ (Roca and Wang, 1994; Roca et al., 1996b; Roca, 2004)) separates. Our data strongly support this concept. Because the hTOP2β core is capable of performing the two critical functions of a type II topoisomerase – supercoil relaxation and DNA decatenation – we hypothesised that an ATPase-less hTOP2β construct (hTOP2β-HL) might complement the temperature-sensitive deficiency of a *top2-4* yeast strain. Interestingly, it did not, nor did an even more catalytically active ATPase-less mutant, hTOP2β-HL^R757W^ (**Fig. 5A**). A potential interpretation of this result is that the level of topoisomerase activity afforded by hTOP2β-HL might be insufficient to provide the essential enzymatic functions that yeast require to grow. However, our data show that the R757W substitution substantially increases the efficiency of strand passage by the headless construct, to the point where its level of activity is either comparable to (supercoil relaxation, **Fig. 3A** vs. **1A**), or within a factor of 2-4 (decatenation, Fig. S5B vs. S2B) from a full-length hTOP2β construct that does support cell viability. Thus, the type IIA topoisomerase core can in the right context be a very robust DNA unlinking enzyme, yet this relatively potent activity does not appear to support yeast cell growth.

In considering why a functional but ATPase-less topo II might be insufficient for cell viability, we noted that the elevated strand passage activities of hTOβP-H2L and hTOP2β-core^R757W^ were both accompanied by an increased damage propensity *in vitro* (**Figs. 2B-C**, **3B-C**). Subsequent genetic studies revealed that cells bearing hTOP2β-HL (either wildtype or the R757W mutant) display a growth defect relative to those with full-length hTOP2β and are inviable in a *rad52* -deficient background (**Fig. 5**). The incompatibility of βh-THOL wPi2th the *rad52-* background demonstrates that the failure of this construct to complement the *top2-4* allele construct is not due to a nuclear localization defect and suggests that it is instead due to cleavage defects introduced into the enzyme; the increased recombination frequency seen upon expression of the hTOP2β headless construct supports this reasoning (**Table S1**). Significantly, appending the ATPase domains back onto the R757W mutant restores much of the integrity of its DNA cleavage-reunion equilibrium *in vitro* (**Fig. 3** v s). anSd4, accordingly, the full-length mutant enzyme can now support cell viability, although it still maintains an increased propensity to damage DNA as seen by the inviability of *rad52*^-^ cells that express the full-length mutant enzyme(Bandak *et al*., 2023). Collectively, these findings establish that the ATPase elements of hTOP2β can mitigate against potent, innate DNA-damaging activities of the hTOP2β c o r *i*e*n vivo*, and additionally support the proposal that type IIA topoisomerases acquired an ATP-binding domain during evolution not to power strand passage *per se*, but to regulate the DNA cleaving activity that accompanies this reaction and suppress DNA damage (Bates, Berger and Maxwell, 2011) (**Fig. 6B**). Future studies will be needed to better understand how the ATPase elements exert this effect at a molecular level.

## Acknowledgements

We thank Lokha R. Alagar Boopathy and Gabriela Swan for assistance with the yeast experiments. This work was supported by the National Institutes of Health [T32-GM008403-28 to A.F.B., R01-CA077373 and R35-CA263778 to J.M.B., CA216010 to J.L.N.] and EMBO [Long-Term Fellowship to T.R.B.].

## Author Contributions

Conceptualization – A.F.B., T.R.B., J.L.N., J.M.B.; Investigation – A.F.B., T.R.B., K.C.N., R.G.; Writing – A.F.B., T.R.B., K.C.N., R.G., J.L.N., J.M.B.; Funding – A.F.B., T.R.B., J.L.N., J.M.B.; Supervision – T.R.B., J.L.N., J.M.B.

## STAR Methods

### Topoisomerase II cloning for protein expression

PCR-amplified full-length topoisomerase genes (hTOP2β: residues 1-1626; hTOP2α: residues 1-1531) were inserted by LIC (ligation-independent cloning (Aslanidis and de Jong, 1990)) into 12UraB (Addgene #48304), a modified version of pRS426 (Christianson *et al*., 1992). The resulting plasmid (hTOP2β-12UraC) encodes a galactose-inducible fusion of the human TOP2B or TOP2A gene with an N-terminal, tobacco etch virus (TEV) protease-cleavable hexahistidine tag. Headless hTOP2β (residues 447-1626) was generated by LIC cloning. Mutant full-length and headless hTOP2β proteins were generated by site-directed mutagenesis of the hTOP2β-12UraC construct using a method based on the QuikChange site-directed mutagenesis protocol. For the topoisomerase core constructs, residues 431-1193 of hTOP2α and residues 447-1206 of hTOP2β were amplified and cloned by LIC into the pET-based vector plasmid 2BT (Addgene #29666), generating IPTG-inducible fusion proteins with an N-terminal, TEV protease-cleavable hexahistidine tag that could be expressed in *E. coli*. Mutant core enzymes were generated by site-directed mutagenesis of the wildtype core constructs using a method based on the QuikChange site-directed mutagenesis protocol.

### Topoisomerase II expression, and purification

Overexpression of full-length and headless (ΔN) constructs were performed in *S. cerevisiae* strain BCY123, with starter cultures grown in complete supplement mixture dropout medium lacking uracil (CSM-URA), supplemented with 2% (vol.vol^-1^) lactic acid and 1.5% (vol.vol^-1^) glycerol as carbon sources. After transformation and growth on CSM-URA+ADE plates at 30 °C, 50 mL CSM-URA starter cultures were inoculoated from single colonies and grown 24h at 30 °C. Starter cultures were transferred to YP expression cultures supplemented with 2% (vol.vol^-1^) lactic acid and 1.5% (vol.vo^-^l^1^) glycerol (100mL starter with 1L media) and grown at 30 °C, 160 rpm, to an OD _600_ of 0.8-1.0 and then induced by the addition of 20 g.l^-1^ galactose. After 6 h incubation with shaking (160 rpm) at 30 °C, cells were harvested by centrifugation (4,500 x g, 15 min, 4 °C), re-suspended in lysis buffer (250 mM NaCl, 1 mM EDTA), and frozen drop-wise in liquid nitrogen.

For purification of proteins expressed in yeast, frozen cells were cryogenically lysed using a Spex 6870 freezer mill, with 15 cycles of 1 min grinding followed by 1 min of cooling. The resultant powder was thawed in A300 (20 mM Tris-HCl [pH 8.5], 300 mM KCl, 20 mM imidazole pH 8.0, 10% [vol.vol^-1^] glycerol with protease inhibitors [1 µg.mL^-1^ pepstatin A, 1 µg.mL^-1^ leupeptin and 1 mM PMSF]) and clarified by centrifugation (17,000 x g, 20 min, 4 °C). The lysate supernatant was passed over an A300-equilibrated HisTrap HP column (GE Healthcare) using a peristaltic pump and washed with 30 mL of A300 and 25 mL of A100 (20 mM Tris-HCl [pH 8.5], 100 mM KCl, 20 mM imidazole pH 8.0, 10% [vol.vol^-1^] glycerol with protease inhibitors). The HisTrap HP column was then connected to an Akta Explorer FPLC (GE Healthcare) and linked upstream of a HiTrap S HP column (GE Healthcare) and equilibrated with a further 5 mL of A100. The tandemly coupled columns were next washed with 25 mL B100 (20 mM Tris-HCl [pH 8.5], 100 mM KCl, 200 mM imidazole pH 8.0, 10% [vol.vol^-1^] glycerol with protease inhibitors) to elute the tagged protein onto the S column, followed by an additional 15 mL of A100 to reduce imidazole levels. A salt gradient was then applied to the coupled columns, reaching 100% buffer C (20 mM Tris-HCl [pH 8.5], 500 mM KCl, 10% [vol.vol^-1^] glycerol with protease inhibitors) over 25 min. Peak fractions were assessed by SDS-PAGE, collected, and concentrated in 100-kDa-cutoff Amicon concentrators (Millipore). His-tagged TEV protease (QB3 MacroLab) was next added to the concentrated samples and incubated at 4 °C overnight. This mixture was then passed over a second HisTrap HP column equilibrated and washed with buffer D (20 mM Tris-HCl [pH 8.5], 500 mM KCl, 20 mM imidazole pH 8.0, 10% [vol.vol^-1^] glycerol) to remove any uncleaved enzyme and the protease. The flowthrough was collected and concentrated, then separated by gel filtration using an S400 column (GE Healthcare) equilibrated in sizing buffer (20 mM Tris-HCl [pH 7.9], 500 mM KCl, 10% [vol.vol^-1^] glycerol). Peak fractions were pooled and concentrated by centrifugation at 4000 RPM using a 30-kDa MWCO filter (Amicon). These final purified samples were then combined with a one-third volume of storage buffer (20 mM Tris-HCl [pH 7.9], 500 mM KCl, 70% [vol.vol^-1^] glycerol), quantified for protein concentration by NanoDrop (ThermoScientific), and snap frozen as 10 µL aliquots for storage at -80 °C.

Wildtype and mutant hTOP2α and hTOP2β core enzymes were overexpressed in *E. coli* strain Rosetta 2 pLysS (EMD Millipore) by growing cells transformed with the appropriate expression vector in 2x YT (at 37 °C and 150 rpm) to an OD _600_ ∼0.3. The temperature was then reduced to 16 °C and cells were grown to an OD _600_ of 0.6-1.0, after which they were then induced with 0.5 mM IPTG and left to grow overnight (20 h) at 16 °C. Induced cells were harvested by centrifugation (4,500 x g, 20 min, 4 °C), re-suspended in buffer A800 (20 mM Tris-HCl [pH 7.9], 800 mM NaCl, 30 mM imidazole pH 8.0, 10% [vol.vol^-1^] glycerol with protease inhibitors), and frozen drop-wise in liquid N _2_. For protein purification, cells were thawed on ice and lysed by 4 cycles of sonication (Misonix Sonicator 3000, 15 sec burst with 2 min rest, on ice). Lysates were clarified by centrifugation (17,000 x g, 30 min, 4 °C) and the supernatant passed over a 5 mL HisTrap HP column (GE Healthcare) equilibrated in A800. Samples were washed in 5 column volumes of A800 and a further 10 column volumes of A400 (20 mM Tris-HCl [pH 7.9], 400 mM NaCl, 30 mM imidazole pH 8.0, 10% [vol.vol^-1^] glycerol with protease inhibitors). Protein was then eluted with B400 (20 mM Tris-HCl [pH 7.9], 400 mM NaCl, 500 mM imidazole pH 8.0, 10% [vol.^-^v^1^]ogl lycerol with protease inhibitors) and concentrated in 30-kDa-cutoff Amicon concentrators (Millipore). 0.5 mg of TEV was added to the concentrated sample, which was then dialysed against 1 L of A400 overnight at 4 °C. This mixture was then passed over a second 5 mL HisTrap HP column (pre-equilibrated with buffer A400) and washed with an additional 5 column volumes of A400. The flowthrough was collected and concentrated in 30-kDa-cutoff Amicon concentrators (Millipore), then separated by gel filtration using an S300 column (GE Healthcare) equilibrated in sizing buffer (50 mM Tris-HCl [pH 7.9], 500 mM KCl, 10% [vol.vol^-1^] glycerol). Peak fractions as determined from SDS-PAGE were pooled and concentrated. The final samples were combined with a one-third volume of storage buffer (20 mM Tris-HCl [pH 7.9], 500 mM KCl, 70% [vol.vol^-1^] glycerol), quantified by NanoDrop (ThermoScientific), and snap frozen as 10 µL aliquots for storage at -80 °C.

### Preparation of plasmid DNA substrates

Negatively supercoiled pSG483 (2927 bp), a pBlueScript SK+ (Agilent) derivative containing a single Nb.BbvCI site, was prepared from *E. coli* XL-1 blue cells (Agilent) using a maxiprep kit (Macherey-Nagel). A portion of this sample was treated with BamHI to form linear plasmid. Another portion of this sample was nicked with Nb.BbvCI and an aliquot was removed to make a nicked pSG483 stock. The kDNA substrate was purchased from Inspiralis; restriction enzymes were from NEB.

### DNA supercoil relaxation and cleavage assays

Prior to use, protein aliquots were thawed on ice for 10 min. The samples were then serially diluted in successive twofold steps using protein dilution buffer (50 mM Tris [pH 7.5], 500 mM KOAc, 2 mM MgOAc, 1 mM TCEP, 50 µ g^-1^. mB SLA and 10% [vol.^-1^v]ogllycerol) to a concentration of 156.25 nM dimer. For drug titrations, a master reaction mixture was made containing four parts diluted enzyme, five parts 4x reaction buffer (40 mM Tris [pH 7.5], 38.4 mM MgOAc, 4 mM TCEP, 100 µg.mL ^-1^ BSA and 32% [vol.vol^-1^] glycerol) and one part of a 500 ng.µL^-1^ solution of substrate (either negatively supercoiled or relaxed) pSG483 plasmid DNA. The mixture was then incubated on ice for 5 min. Drug titrations were prepared by mixing 2 µL of an appropriate drug dilution (or 2 µL of solvent for “zero drug” controls) with 1 µL of 20 mM ATP and 7 µL of ddH _2_O. These 10-µL drug mixtures were then added to 10-µL aliquots of the reaction mixture on ice, quickly transferred to 37 °C, and incubated for 30 min. Final reaction conditions consisted of 31.25 nM full-length dimers, 12.5 nM supercoiled pSG483, variable drug content (or solvent), 1 mM ATP, 20 mM Tris [pH 7.5], 100 mM KOAc, 10 mM MgOAc, 1.2 mM TCEP, 35 µg.mL^-1^ B S A a n d 1 0 % [^-1^v] golylc.ervool. Fl ollowing incubation, the reactions were quenched with 2 µL of stopping buffer containing either 5% (wt.vol^-1^) SDS (for etoposide-containing reactions and reversible cleavage measurements) or 5% (wt.vol^-1^) SDS and 125 mM EDTA (for irreversible cleavage measurements). Stopped reactions were subsequently treated with 1 µL of 12 mg.mL^-1^ proteinase K, followed by further incubation at 37 °C for 30 min.

Reactions were then stored on ice until immediately before gel loading, whereupon a 6x agarose gel loading dye was added to the samples and the solutions warmed to 37 °C for 5 min. Supercoil relaxation samples were separated by electrophoresis in 1.4% (wt.vol^-1^) TAE agarose gels (50 mM Tris-HCl [pH 7.9], 40 mM NaOAc, and 1 mM EDTA [pH 8.0] running buffer), for 6–15 h at 2–2.5 V.cm ^-1^.; for cleavage assays, conditions were the same except that XXX [conc.] of ethidium bromide was included with the gel (not the running buffer). To visualize the DNA, native gels were poststained with 0.5 μg.mL^-1^ ethidium bromide in TAE buffer for 30 min, destained in TAE buffer for a further 30 min, and exposed to UV illumination. Supercoiled relaxation/cleavage assays performed with the core enzymes followed the protocol detailed above, except that final concentration of core dimers used in the assays was 125 nM. For enzyme concentration or DMSO titration studies, the same protocol was also followed, except that different amounts of protein or DMSO were included in final reaction volumes as described in the text and figure legends. Gel images were analyzed using ImageJ (Schneider, Rasband and Eliceiri, 2012), and data were plotted using Prism (GraphPad Software).

### Yeast complementation

Previous topoisomerase complementation work has relied on regulatable vectors (based on the Gal1/10 promoter) or constitutive expression with vectors that included yeast sequences at the amino terminus (Jensen *et al*., 1996; Meczes *et al*., 1997). We constructed a single copy vector (pKN17) driving full-length hTOP2β (with no yeast coding sequences) from the yeast TPI promoter, derived from pRS112. This vector efficiently complements a *S. cerevisiae top2-4* strain, allowing for growth at a non-permissive temperature (**Fig. 5A**).

### Construction of variant Top2β expression constructs for examination of yeast phenotypes

The plasmid pKN17 was used as a template to generate the PCR products for cloning hTOP2β amino acids 449-1626 construct (hTOP2β-HL) using standard Gibson assembly protocols (Gibson *et al*., 2009). The first set of primers amplify the segment containing part of URA3 until the end of the TPI promoter into part of the truncated hTOP2β sequence (underlined). The start codon (ATG) is in bold (Forward-5’-*AACACATGTGGATATCTTGACTGATTTTTCCATGGAGG* -3’ and Reverse-5’-<u>TTACTGATGA**CAT**</u>GGGGCTGCAGGAATTCCTG-3’). The second set of primers generate a product containing hTOP2β (underlined) starting from serine 449 until the URA3 marker in YCplac33. URA3 sequences are in italics (Forward-5’-TTCCTGCAGCCCC<u>**ATG**TCATCAGTAAAATACAGTAAAATC</u> -3’ and Reverse-5’-*AATCAGTCAAGATATCCACATGTGTTTTTAGTAAAC* -3’). PCR products were set up using iProof High-Fidelity PCR kit and the GC buffer (Bio-Rad). The assembled reactions were transformed into NEB 5-alpha high efficiency competent *E. coli*. An MfeI/BamHI fragment containing the hTOP2β R757W mutation was removed from the hTOP2β-12UraC R757W construct and ligated into the MfeI/BamHI digested hTOP2β headless. A full list of primers is available (**Table S2**).

### Assessment of rad52-dependent lethality

Plasmids were transformed into the isogenic strains JN362a (*RAD52*+) and JN394 (*rad52*-) strains using lithium acetate. Transformants were plated to ura-agar plates and incubated 3-5 days at 30 °C. Strains expressing wildtype or mutant hTOP2β typically required 1-2 days more for colony formation than cells transformed with an empty vector or expressing other eukaryotic topoisomerases. Plates were photographed immediately after removal from the incubator.

## Supplemental Figure Legends

**Fig. S1.** DMSO enhances supercoil relaxation activity of the hTOP2α and hTOP2β cores. (A) Activity of the hTOP2α and hTOP2β cores on negatively supercoiled plasmid DNA as titrated against increasing concentrations of DMSO. (B) Substitution of the catalytic tyrosine in the hTOP2α and hTOP2β cores abolishes ATP-independent supercoil relaxation activity.

**Fig. S2.** The hTOP2β^R757W^ core has enhanced decatenation activity. (A) Location of Arg757 (PDB: 3QX3). Decatenation of kDNA by the wildtype hTOP2β core, the hTOP2β^R757W^ core and wildtype, full-length hTOP2β. Increasing amounts of each enzyme were incubated with 250 ng of kDNA for 30 min at 37 °C, in the absence ((-), for the cores) or presence ((+), for full-length) of ATP.

**Fig. S3**. The hTOP2β^K600T^ core has enhanced supercoil activity and cleavage propensity relative to the wildtype hTOP2β core. Ratios refer to enzyme dimers:DNA. Enzymes were incubated with 10 ng/μL of negatively supercoiled pSG483 for 30 min at 37 °C.

**Fig. S4**. Full-length hTOP2β^R757W^ has reduced cleavage activity compared to the hTOP2β^R757W^ core. Activity of full-length hTOP2β^R757W^ on negatively supercoiled plasmid DNA, separated in agarose gels in the absence (upper panel) or presence (lower panel) of ethidium bromide (EtBr). Compare with **Fig. 3**.

**Fig. S5**. Headless hTOP2β^R757W^ has enhanced supercoiling and decatenation activities over wildtype headless hTOP2β. (A) Activity of headless wildtype hTOP2β and hTOP2β^R757W^ on negatively supercoiled plasmid DNA. Ratios refer to enzyme dimers:DNA. (B) Decatenation of kDNA by headless wildtype hTOP2β and hTOP2β^R757W^. Increasing amounts of each enzyme were incubated with 250 ng of kDNA for 30 min at 37 °C.

## Supplemental Tables

**Table S1.**
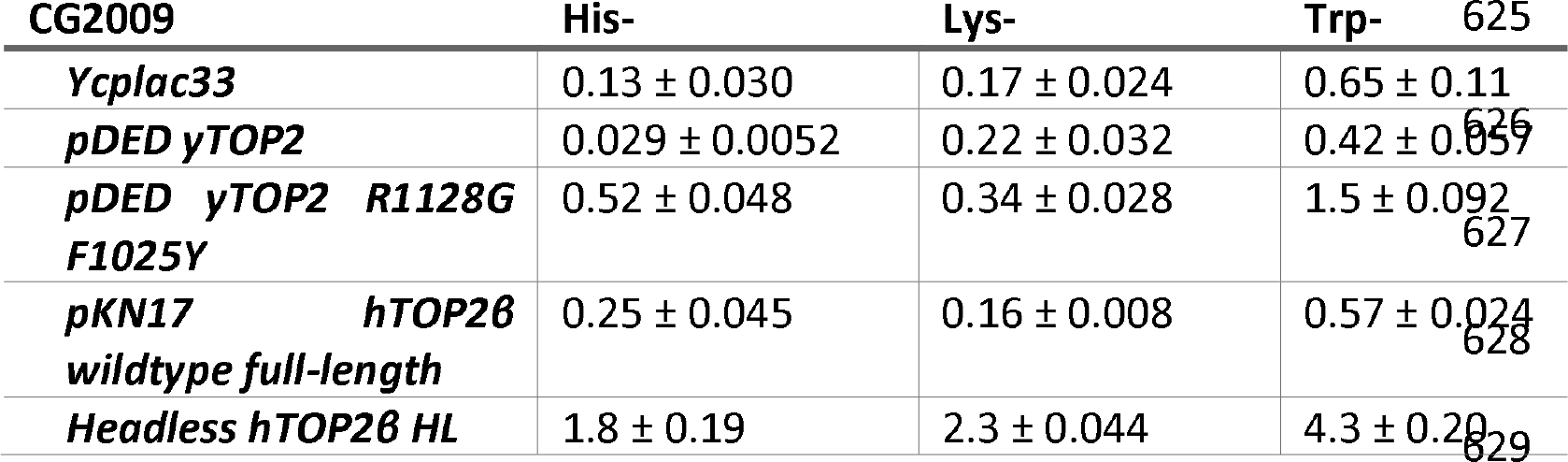
Recombination frequency induced by expression of headless hTOP2β HL. CG2009, genotype MATa/MATa lys2-1/lys2-2 tyr1-1/tyr1-2 his7-2/his7-1 leu2D/leu2D ura3D/ura3-1 trp5-d/trp5-c met13-d/met13-c ade5/ADE5 ade2/ade2 was transformed with the indicated plasmids, and individual single colony transformants were inoculated in yeast synthetic complete media. After overnight incubation, cultures were plated to media lacking histidine (His-) lysine (Lys-) or tryptophan (Trp-) to select for heteroallelic recombination at HIS7, LYS2 or TRP5. Diluted cultures were also plated to media lacking uracil to determine the number of viable plasmid carrying cells. Recombination frequencies were calculated by dividing the number of colonies on the relevant plates by the total viable count. Recombination frequencies are expressed x10 ^4^ and are shown mean ± SD from five independent transformants. pDEDyTop2 overexpresses wildtype yTOP2 from the DED1 promoter. pDED yTOP2 R1128G F1025Y expresses a self-poisoning TOP2 allele that leads to elevated recombination and is a positive control for the recombination induction(Stantial *et al*., 2020).

**Table S2.**
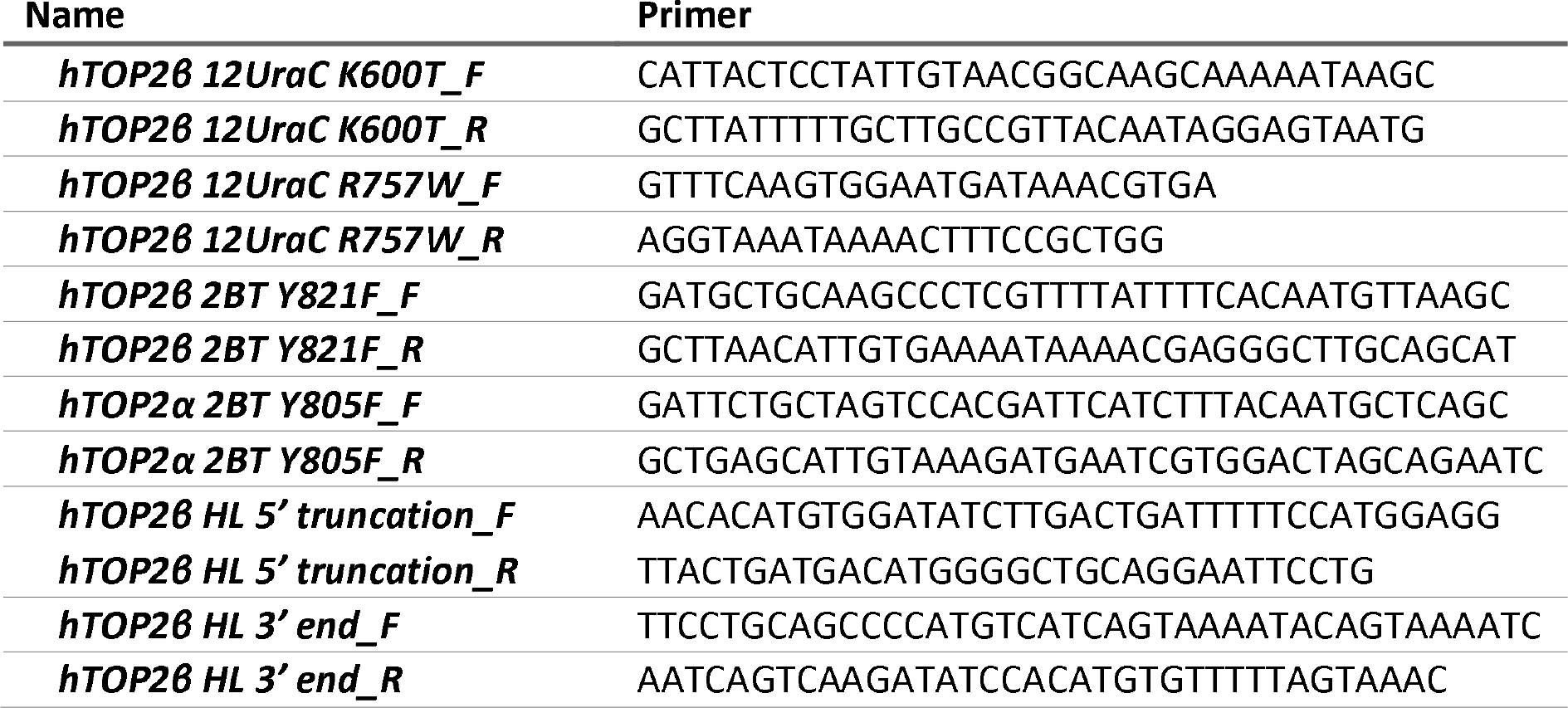
Primers used in this study.

