## Supplementary figures and images for "Using energy to go downhill – a genoprotective role for ATPase activity in DNA topoisomerase II"

### Supplemental Figures

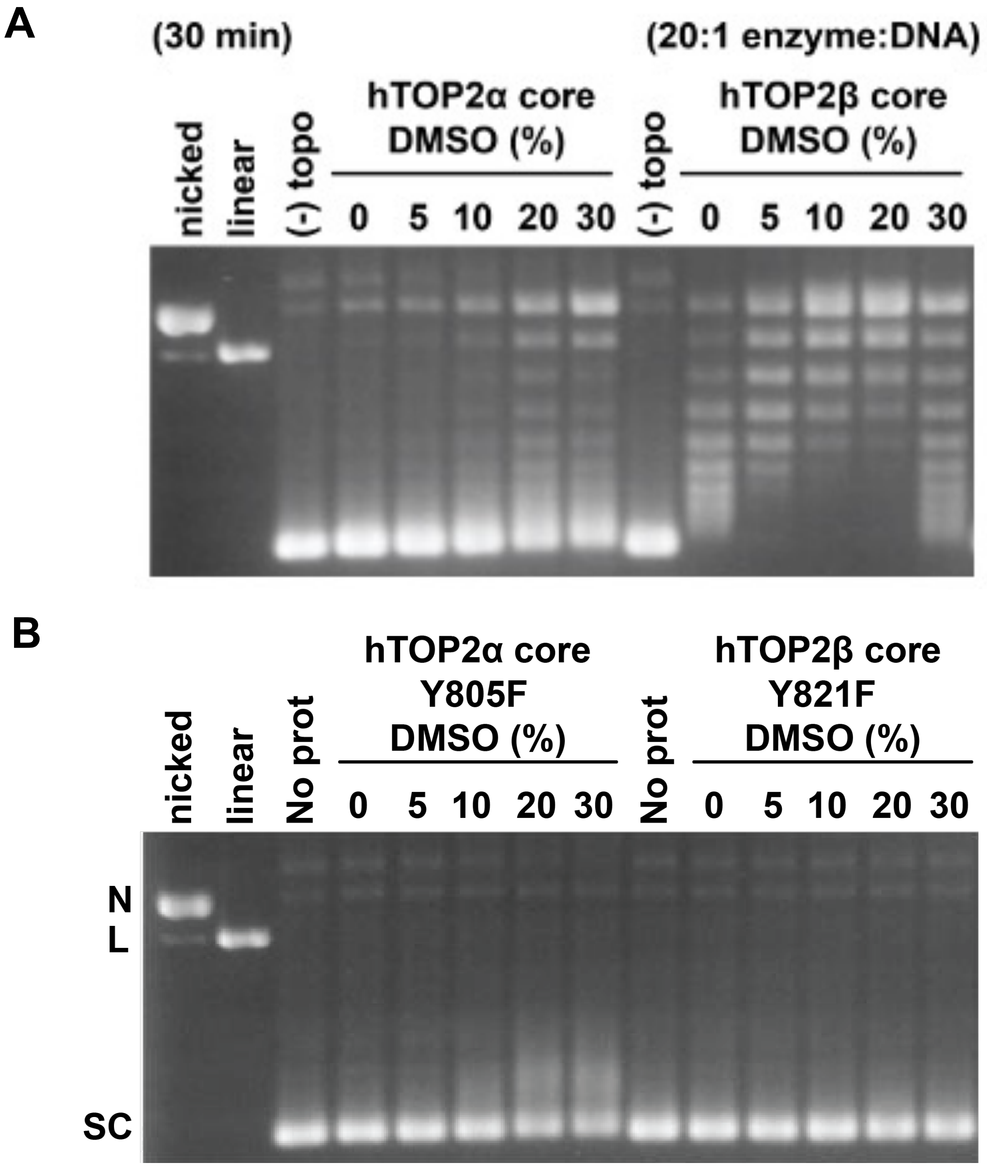

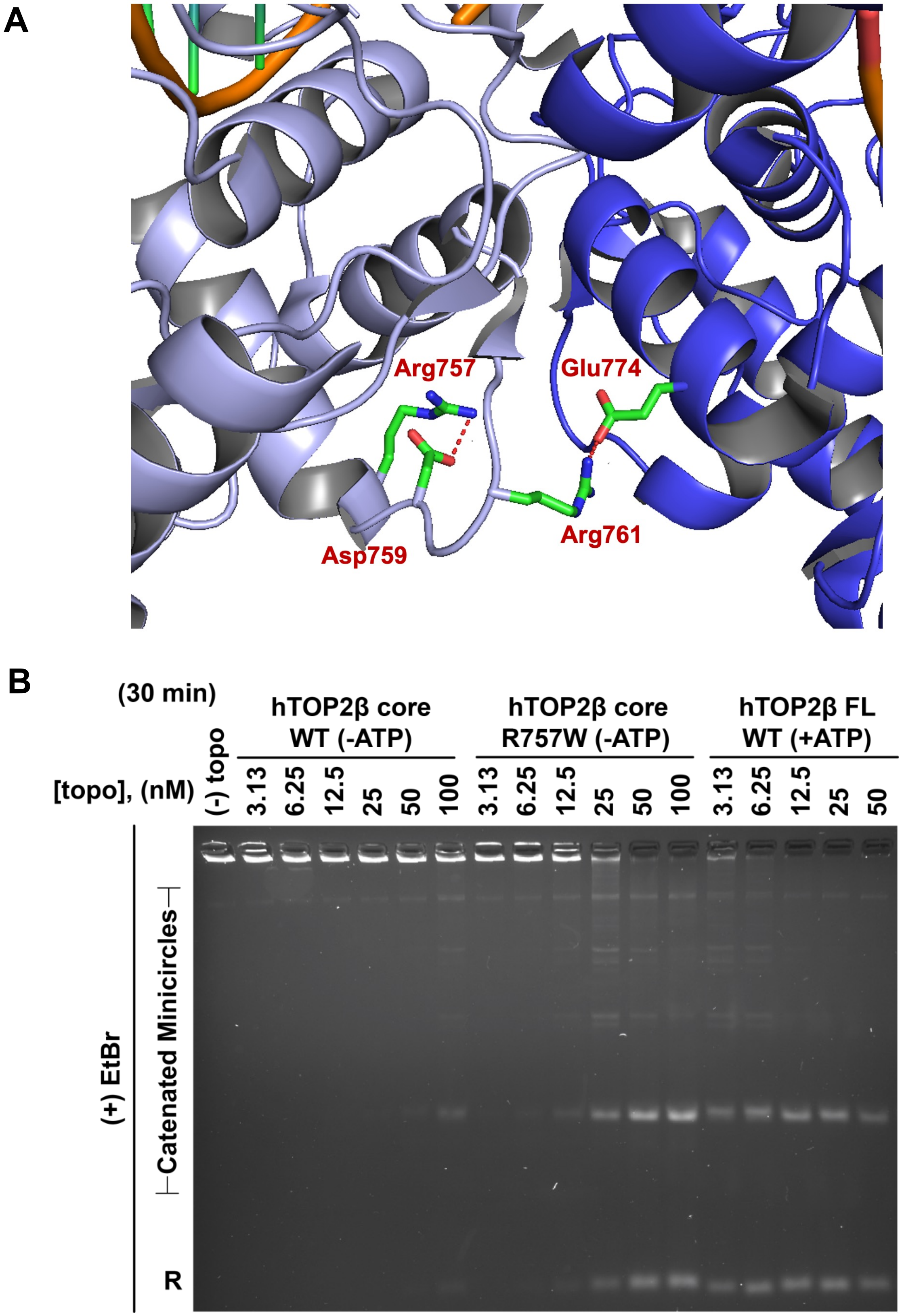

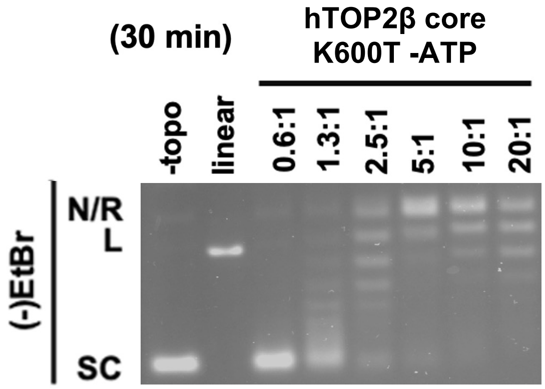

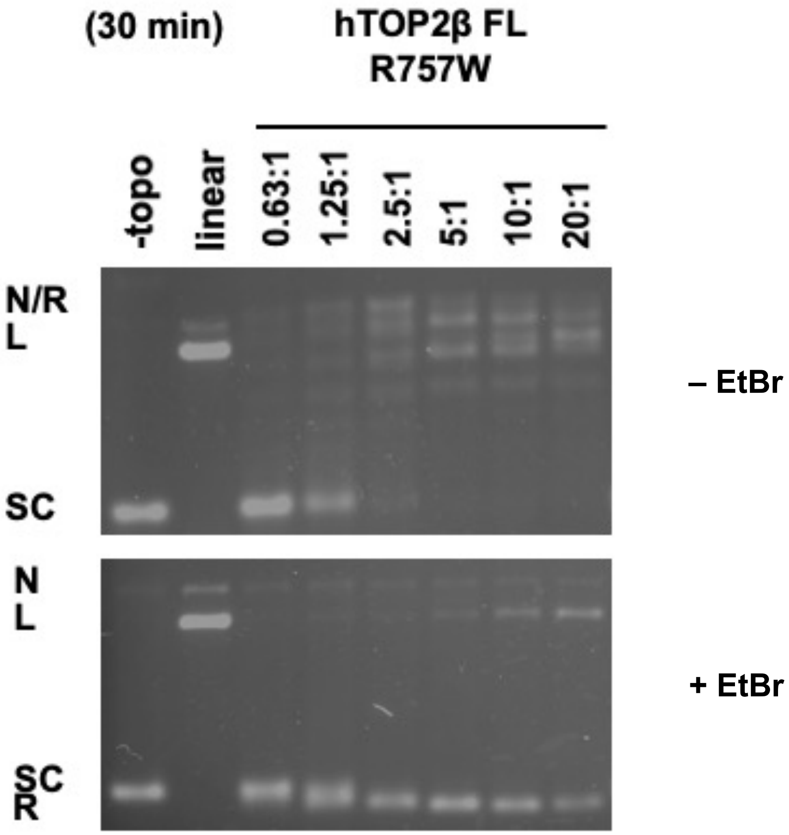

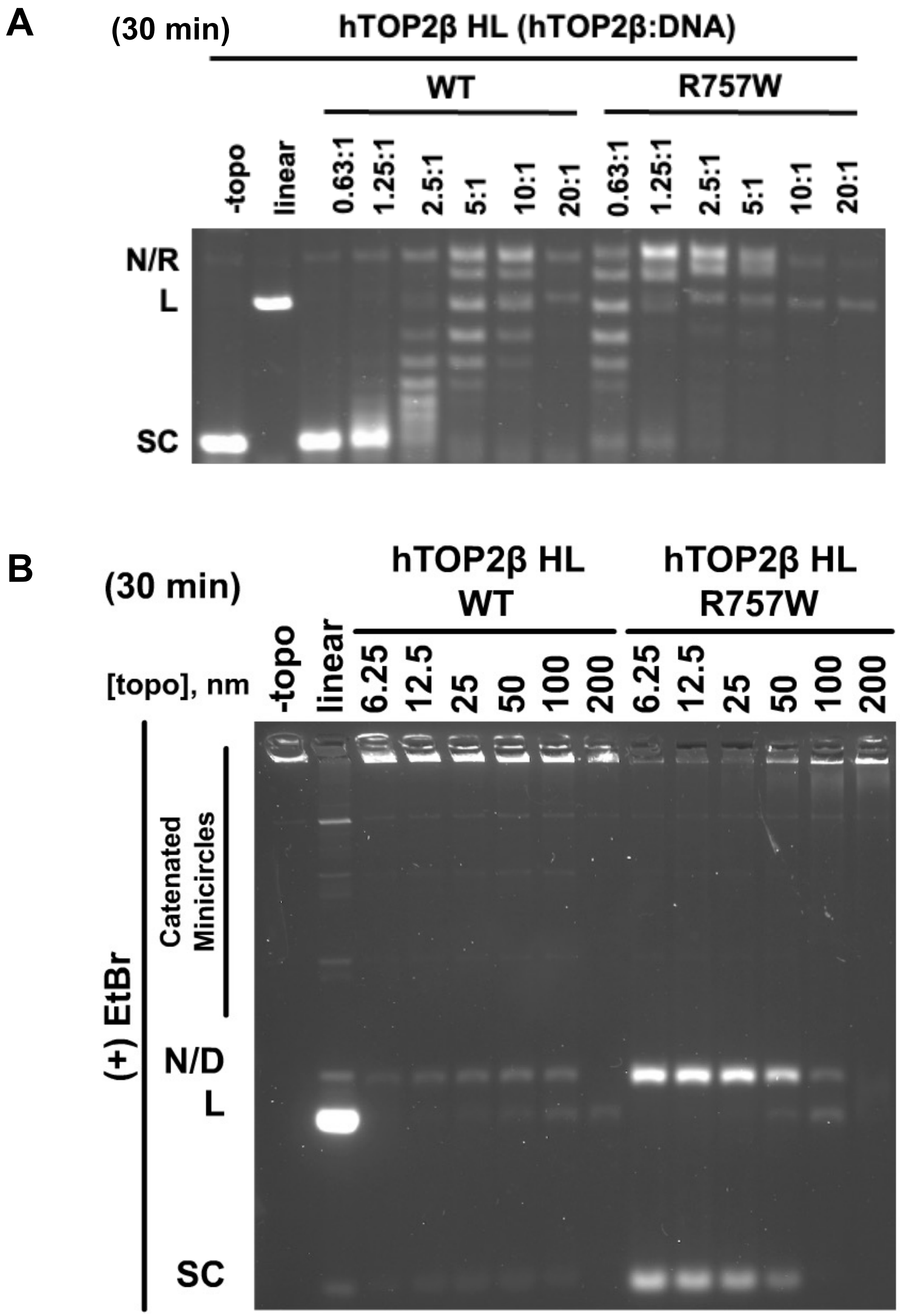
